# Bioengineered human colon organoids with *in vivo*-like complexity and function

**DOI:** 10.1101/2023.10.05.560991

**Authors:** Olga Mitrofanova, Nicolas Broguiere, Mikhail Nikolaev, Matthias P. Lutolf

## Abstract

Organoids and microphysiological systems, such as organs-on-a-chip, have emerged as powerful tools for modeling human gut physiology and disease *in vitro*. However, although physiologically relevant, these systems often lack the environmental milieu, spatial organization, cell-type diversity, and maturity necessary for mimicking adult human intestinal mucosa. To instead generate models closely resembling the *in vivo* cell-type composition and spatial compartmentalization, we herein integrated organoid and organ-on-a-chip technology to develop a primary human stem–cell-derived organoid model, called ‘mini-colons’. The luminal access and flow in human mini-colons removes shed cells to greatly enhance tissue longevity and differentiation over physically inaccessible human intestinal organoids that accumulate trapped cellular debris and waste. By establishing a gradient of growth factors, we replicated and sustained *in vivo*-like cell fate patterning and concurrent differentiation to secretory cell types and colonocytes. These long-lived human mini-colons contain abundant mucus-producing Goblet cells that lubricate the colonic epithelial lining. The stem and proliferative progenitor cells are also realistically confined to the crypts, facilitating stable homeostatic tissue turnover and preserving tissue integrity for several weeks. Also signifying mini-colon *in vivo*-like maturation, single-cell RNA sequencing showed emerging mature colonocytes and absorptive BEST4^+^ colonocytes. This methodology could be expanded to generate microtissues derived from the small intestine and incorporate additional microenvironmental components, thus emulating the intricate complexity of the native gut in an *in vitro* setting. Our bioengineered human organoids provide a highly accurate, long-lived, functional platform to systematically study human gut physiology and pathology, and for the development of novel therapeutic strategies.

## Introduction

Human intestinal organoids offer remarkable prospects for modeling intestinal homeostasis and disease. Although it is now possible to recreate many physiological features of the human gut *in vitro*, current approaches to generate organoids produce closed cystic structures with limited cell-type variability, hampering their relevance and applicability^1^. While murine intestinal organoids effectively balance stem-cell maintenance and differentiation in crypt-like and villi-like domains^2^, respectively, human intestinal organoids have not yet achieved the same level of homeostasis and spatial organization^3,4^. Recent research on developing fetal human intestines^5,6^ and the discovery of factors that regulate differentiation programs of distinct intestinal cell lineages^7^ have informed advances in intestinal organoid culture. However, a fully accurate biomimetic niche for human colonic and small intestinal organoid growth *in vitro* is still missing.

Human intestines-on-a-chip have emerged as an alternative approach for mimicking gut epithelia *in vitro*. In contrast to organoids, these platforms contain microfluidic channels that create dynamic systems with realistic fluid flow, thus allowing greater experimental control of the developing tissues. Intestines-on-a-chip typically incorporate hollow channels that are colonized by intestinal epithelial cells to produce flat planar cultures that bear a limited resemblance to the anatomical cytoarchitecture of the native gut^8–12^. However, approaches that utilize 3D scaffolds to preserve crypt-villus features rely on supportive membranes such as Transwell cell culture inserts that prevent co-culturing with other non-epithelial cell types and overcomplicate experimental tractability and image-based readouts^13–15^. While these models recapitulate some aspects of the native human gut, it has been challenging to achieve cell-type variability and long-term homeostasis.

Recently, we overcame these limitations by combining microfabrication, hydrogel engineering, and stem-cell intrinsic self-organization to guide the morphogenesis of mouse Lgr5^+^ intestinal stem cells into tubular and perfusable intestinal tissues with characteristic crypt and villus domains similar to native intestine^16^. Considering the lack of advanced *in vitro* models accurately representing human intestinal physiology, we reasoned that extending this approach to using patient primary cells and leveraging recent advances in organoid culture, would allow to produce more *in vivo*-relevant colonic and small intestinal tissues with enhanced cellular diversity, longevity, and function. To this end, we generated human mini-colons and mini-intestines from primary stem cells and demonstrated their close functional and morphological resemblance to the native counterparts and amenability to a multitude of biological and translational applications.

## Results

### Bioengineering human mini-colons

To create human mini-colon tubes capable of supporting fluid flow, we employed and improved upon a previously introduced hybrid organ-on-a-chip system that incorporates a central hydrogel scaffold within a polydimethylsiloxane (PDMS) device^16^ (**Figures 1A and 1B**). The biomimetic hydrogel was modified with laser ablation to generate a microchannel approximating the topography and dimensions of native human colon (**Figure 1B)**. The crypts were then populated with human colon cells, which attached rapidly over the next days to form a coherent and tightly packed epithelium that thickened over time (**Figures 1C and 1D**). Mini-colons were successfully generated using three different donors, and all exhibited consistent rates of microchannel epithelialization (**Figure 1D**). In all mini-colons, the resulting epithelium formed a contiguous tube-like monolayer that expressed high levels of E-cadherin at the cell-cell junctions and developed outward-projecting invaginations of the deep crypts (**Figure 1E, Supplementary video 1**). A histological analysis confirmed that mini-colons remained open on both ends, forming a hollow epithelial tube lined by columnar cells with basally oriented nuclei (**Figure 1F**). We next assessed the mini-colons for barrier integrity by perfusing a fluorescent probe (40 kDa dextran) through the lumen, which revealed that fully epithelialized tubular mini-colons established an intact leak-tight barrier under homeostatic conditions (**Figure 1G**).

**Figure 1.**
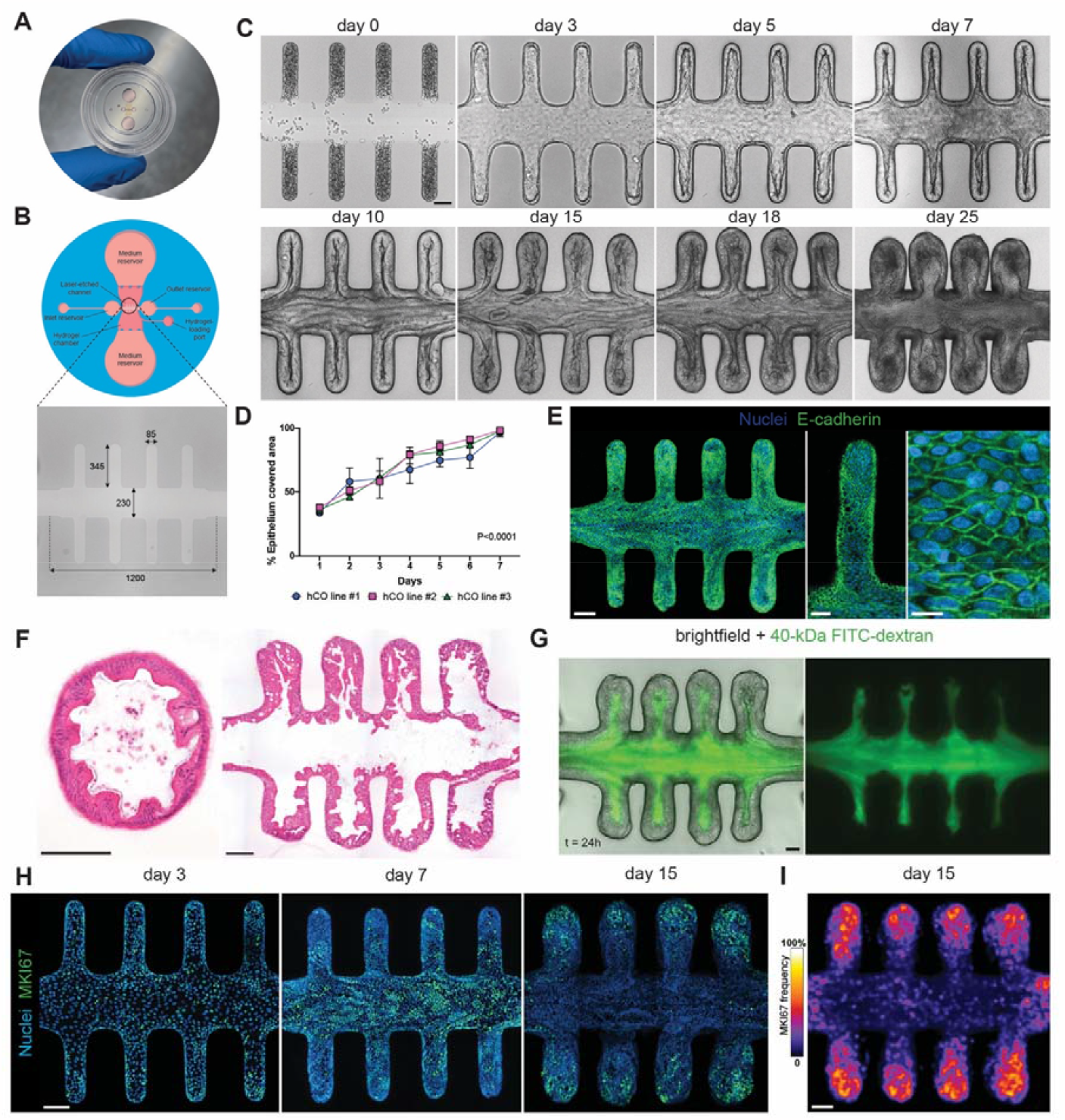
Bioengineered human mini-colons establish homeostatic culture of colonic epithelium. **A**, A photograph of the microfluidic chip used in the study. **B**, A schematic overview of the modified hybrid organ-on-chip incorporating central hydrogel chamber within a PDMS device and dimensions (in μm) of an open microchannel with an *in vivo*-like anatomical structure, generated by laser ablation within a central hydrogel compartment. **C**, Brightfield time-course experiment of the epithelium formation in tissue-engineered human mini-colons. **D**, Quantification of the epithelium coverage area over the course of 7 days. The epithelium area was segmented using machine learning toolkit ilastik and quantified using custom-made ImageJ plugin. The data are shown for 3 independent organoid lines which were used at passage 6-10. **E**, Fluorescence confocal image of a 10-day old epithelial tube stained for tight junction protein E-cadherin (green) and DAPI (blue, nuclei). **F**, Histological transverse and longitudinal cross-sections of 20-day-old mini-gut tubes stained with H&E. **G**, Brightfield and fluorescence confocal images of a 15-day-old mini-colon perfused with fluorescein isothiocyanate (FITC)-tagged dextran (40 kDa), showing the maintenance of epithelium integrity. **H**, Fluorescence images showing localization of MKI67^+^ proliferative cells in mini-colons at different time points. **I**, Frequency map showing the localization of MKI67^+^ proliferative cells in 15-day-old tissues. Average of the maximal intensity projection of a *z*-stack of around 150 μm for *n* = 8 mini-colons. Scale bars = 100 μm, magnification = 50 μm.

### Establishing a long-term homeostatic culture of human mini-colons

Reproducing rapid intestinal cell turnover and multilineage differentiation in human intestinal organoids has been challenging. Previous studies demonstrated that mature differentiated cell types cannot emerge in human small intestinal organoids before the mitogenic factors are removed, but this comes with a significant drawback of preventing continued organoid expansion^4^. To enable concurrent multilineage differentiation and self-renewal in human small intestinal organoids, Sato and colleagues replaced the p38 inhibitor with insulin-like growth factor 1 (IGF1) and fibroblast growth factor 2 (FGF2)^3^. Nevertheless, mature absorptive cell types are largely underrepresented in human intestinal organoids^3^ and require the activation of BMP signaling, which also drives the characteristic zonation along the crypt-villus axis in the small intestine^18^. However, the coordination of cell-type composition within colonic tissues and the ability of human colon organoids to accurately replicate organomimetic cell-type diversity have yet to be investigated.

Here, we hypothesized that because the mini-colon platform allows access to both the apical and basal sides, we could apply signaling gradients that would aid in both mature differentiation and self-renewal to generate longer lasting and more biologically relevant cultures. More specifically, we could integrate previously established protocols to induce secretory differentiation and self-renewal from the basal side while simultaneously promoting apical differentiation towards colonocytes. Strikingly, exposing the lumen to the differentiation medium after one week of culture resulted in epithelial cell shedding, and removing these accumulating shed cells necessitated repeated perfusions applied every 24 hours (**Figure S1A, Supplementary video 1**). Using a rocking platform culture, we effectively extruded a mucus-encapsulated mass of shed dead cells from the lumen (**Figure S1B**). Overall, integrating perfusion and luminal flow dramatically augmented growth and tissue longevity, allowing us to establish a one–month-old culture of human mini-colon tubes while preserving the overall anatomy of the tissue (**Figure 1C**). Moreover, this approach both replicates the homeostatic self-renewal of epithelial lining in human mini-colons and simulates the biomimetic patterning of proliferative progenitor cells. For instance, during the early stages of epithelium formation, human mini-colons were initially populated by highly proliferative cells distributed throughout the tissue. Upon switching to differentiation, these cells gradually become restricted to the crypt regions, creating proliferative zones similar to *in vivo* (**Figures 1H and 1I**).

### Cellular diversity of human mini-colons

This crypt-restricted localization of proliferative cells prompted us to further investigate whether differentiation-promoting conditions could also produce the stereotypical patterning of the key colonic cell types. To interrogate cellular diversity, we conducted multiplexed scRNA-seq using cell hashing^19^ of conventional human mini-colons in biological triplicates at day 14 (1 week of differentiation) and day 21 (2 weeks of differentiation) of culture. Mini-colon cells were classified via unsupervised clustering into 7 discrete subpopulations, which were further visualized with uniform manifold approximation and projection (UMAP) (**Figure 2A**). We annotated each cell population using canonical genes, which identified that human mini-colons harbor most of the cell types of the native colon, including a population of dividing cells; stem and progenitor cells; mature colonocytes, including crypt-top colonocytes and newly identified absorptive BEST4^+^ colonocytes; as well as Goblet and enteroendocrine cells (**Figures 2A and 2B)**. In addition, scoring and projecting the cell-cycle state on the UMAP plot revealed a clustering of cells in S and G1 phases, corresponding to stem cells and dividing transit-amplifying (TA) cells (**Figure S1C**).

**Figure 2.**
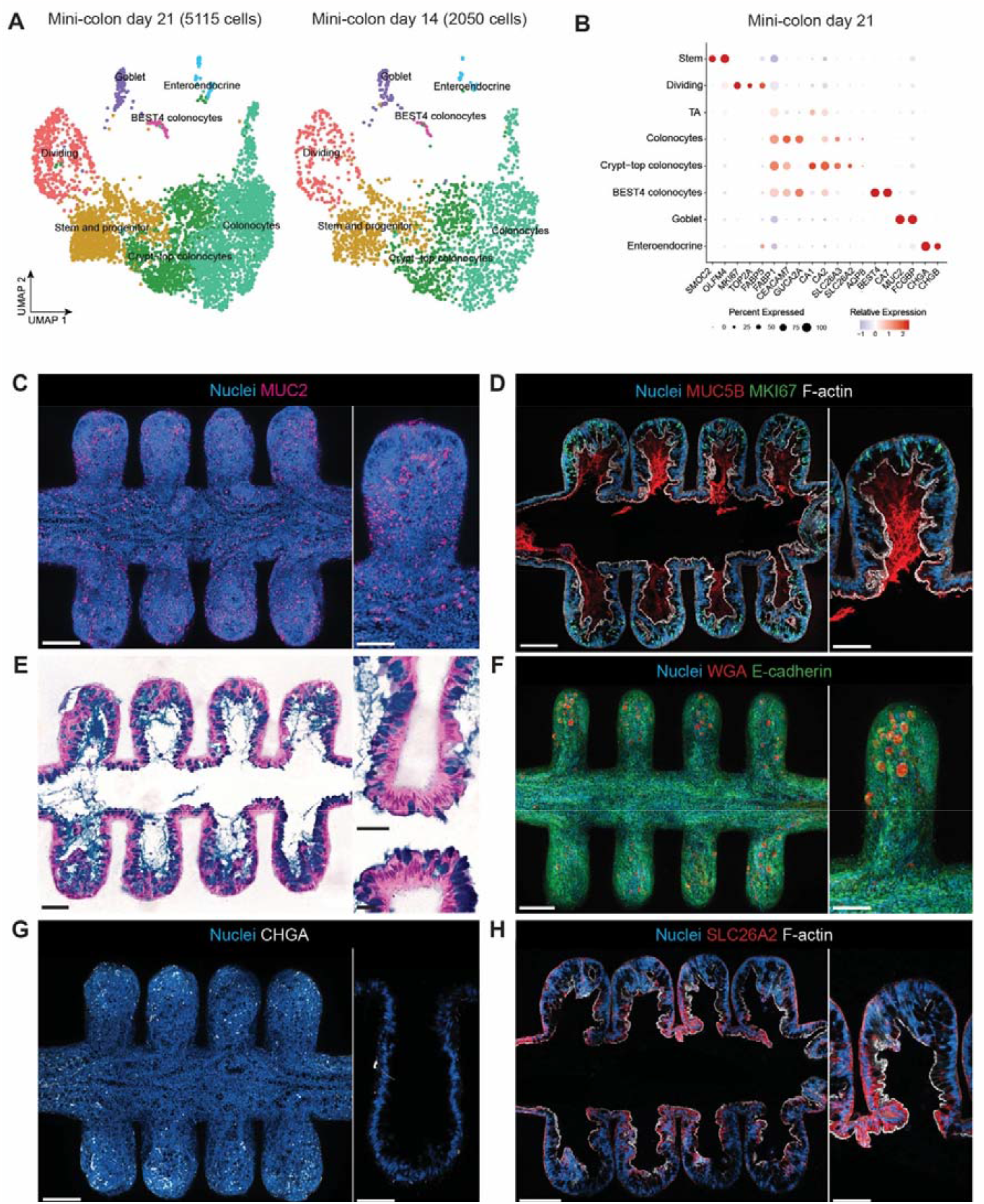
Human mini-colons capture colonic cellular diversity and cell type patterning. **A**, UMAP of the key intestinal epithelial cell types in 21-day-old mini-colons (n = 5115 cells) and 14-day-old mini-colons (n = 2050 cells). **B**, Dot plot showing the relative expression and the percentage of cells expressing selected markers across scRNA-seq clusters of 21-day-old mini-colons. **C**, Immunofluorescence image of human mini-colon histological section stained for Goblet cell marker MUC2 (pink) and DAPI (blue, nuclei). **D**, Immunofluorescence image of human mini-colon histological section stained for mucus MUC5B (red), MKI67^+^ proliferative cells (green) and F-actin (white). **E**, Histological longitudinal cross-sections of 18-day-old mini-gut tubes stained with Alcian Blue showing intracellular granules filled with mucus and acidic polysaccharides of the mucus layer (blue) counterstained with Nuclear Fast Red. **F**, WGA (red) lectin staining of the fixed human mini-colon co-stained with DAPI (blue, nuclei) and E-cadherin (green). **G**, Immunofluorescence image of human mini-colon histological section stained for EEC marker CHGA (white) and DAPI (blue, nuclei). **H**, Immunofluorescence image of human mini-colon histological section stained for colonocyte marker SLC26A2 (red), DAPI (blue, nuclei) and F-actin (white). Scale bars = 100 μm, magnification = 50 μm.

Individual replicates of these scRNA-seq studies demonstrated a high degree of uniformity, highlighting the consistency of the experimental procedures and reproducibility of our system (**Figures S1D and S1E**). We found that mini-colons contained a large proportion of mature colonocytes (around 40%), which *in vivo* play critical roles in absorbing water and short-chain fatty acids^20,21^, and shape gut microbiota, by mediating a switch between homeostatic and dysbiotic communities^22^ (**Figure S1E**). Moreover, mini-colons comprised a small proportion of BEST4 colonocytes (from 0.6% on day 14 to 1% on day 21). Additionally, day 21 mini-colons produced more enteroendocrine cells (1.7% vs. 0.9% on day 14). Overall, we did not observe major shifts in cell type composition between day 14 and day 21 mini-colons, with proportions of stem and progenitor (around 25%), dividing (around 15%), and Goblet (around 3%) cells remaining constant, suggesting that upon a week of differentiation, mini-colons achieve a state of homeostasis and maintain a stable ratio of differentiated and proliferative cell types (**Figure S1E**).

### Mucus production by Goblet cells in human mini-colons

To corroborate the scRNA-seq findings, we performed an immunofluorescence analysis, which revealed abundant MUC2^+^ Goblet cells distributed throughout the mini-colons, though primarily localized towards the crypts (**Figure 2C**). However, scRNA-seq revealed the presence of only a small fraction of Goblet cells (**Figure S1E**), suggesting that these cells may be lost during tissue dissociation. In the colon, secreted MUC2 protein together with other components form a thick mucus layer that acts as a protective barrier against microorganisms while allowing the passage of small molecules^24^. To investigate properties of the mucus layer, *ex vivo* experiments with explant tissues or *in vivo* experiments using animal models are often performed^25^. However, the scarcity of human-relevant *in vitro* models^11,26,27^ largely hinders our understanding of the dynamics of the mucus barrier in physiological and pathophysiological conditions in the colon. Moreover, there is no model with preserved 3D cytoarchitecture for investigating how a stem cell niche is shielded by mucus.

Our model may be the first that can fill this gap. Starting from day 5 of our culture, we observed an accumulation of thick transparent mucus that could be collected by accessing the apical side of mini-colons (**Supplementary video 2**). While MUC2 is the major gel-forming mucin in the colon, other mucin proteins also contribute to mucus formation^28^. As such, the mini-colons also contained MUC5B, which is abundantly expressed in the crypt base Goblet cells *in vivo*^23^. Immunostaining revealed MUC5B-stained mucus plumes emanating from the bottom of the crypt (**Figure 2D**). Acidic mucopolysaccharides stained with Alcian blue identified mucin-filled vesicles in Goblet cells and a thick mucus layer covering the apical side of the epithelium. Notably, we captured a continuous release of mucus from Goblet cells, characterized by the discharge of Alcian–blue-stained intracellular granules in histological sections of mini-colons (**Figure 2E**). Strikingly, without perfusion, the lumen of the mini-colons filled with mucus (**Figure S1F**). Although Goblet cells are considered the major intestinal mucus producers, we also detected the expression of MUC5B and membrane-bound mucins MUC1, MUC12, MUC13 and MUC20 in colonocytes, consistent with previous findings^23^ (**Figure S1G**). Further confirming these data, wheat germ agglutinin (WGA) lectin staining detected the presence of glycosylated mucin vesicles within crypt-residing Goblet cells in fixed samples of mini-colons (**Figure 2F, Supplementary video 2**). Overall, human mini-colons provide a unique platform to study mucus secretion with unprecedented detail and tractability. By enabling the visualization and manipulation of the mucus layer *in vitro*, this system has the potential to uncover new insights into the complex interplay between mucus, the microbiota, and the host and to identify novel therapeutic targets for a range of gastrointestinal diseases.

### Diversity of enteroendocrine cells and mature colonocytes in human mini-colons

The immunofluorescence analysis also revealed the presence of numerous CHGA^+^ enteroendocrine cells (EECs) in mini-colons (**Figure 2G**). These EECs were classified based on hormone expression into subtypes of EEC progenitor cells expressing NEUROG3, hormone-secreting populations of L-cells (GCG^+^ and PYY^+^), and enterochromaffin cells (TPH1^+^), which have been previously characterized by scRNA-seq of human colonic biopsies^23^ (**Figure S1H**). The scRNA-seq identified a vast heterogeneity of colonocytes in human mini-colons that has not been comprehensively described in any *in vitro* system (**Figures 2A and 2B**). We thus reconstructed the colonocyte lineage trajectory using diffusion pseudotime analysis to reveal a common differentiation path originating from stem and progenitor cells leading to early colonocytes and then diverging into two lineages of late and BEST4^+^ colonocytes (**Figure S1I**). We also characterized the colonocytes in human mini-colons using the expression of typical *in vivo* colonocyte markers such as CEACAM7 and GUCA2A^23,29^. Furthermore, we also distinguished a subpopulation of crypt-top colonocytes, marked by the expression of markers AQP8, MS4A12, SLC26A2, SLC26A3, CA1, and CA2^23,29^, commonly observed in mature late colonocytes *in vivo* (**Figures 2B and S1J**). Various ion transporters also exhibit distinct regional differences *in vivo*, with a particular abundance of SLC26A2 in colonocytes^23^. Interestingly, immunostaining of human mini-colons revealed SLC26A2-positive cells primarily localized to the flat tops of the crypts (**Figure 2H**), which is consistent with their spatial arrangement *in vivo*^21^. Recently identified BEST4^+^ absorptive colonocytes^29^ were also present in human mini-colons and showed expression of colon-specific OTOP2 and CA7 markers (**Figure S1J)**. This diversity of mature colonocytes reflects a high level of tissue maturity of human mini-colons, which offers a unique platform to functionally investigate colon-specific physiological roles. Overall, scRNA-seq and an immunofluorescence analysis collectively demonstrated that human mini-colons cultured with advanced microenvironmental conditions could closely recapitulate the cellular diversity of the native colon.

### Functionality of human mini-colons

We next studied the ability of our mini-colons to maintain the absorptive function of native colonic mucosa. Although the primary role of the colon is to generate a dense mucus layer that acts as a protective barrier and a habitat for enteric bacteria, it also performs a pivotal task in absorbing residual water and nutrients, particularly short-chain fatty acids^21,30^. As such, choline transporters SLC44A1 and SLC44A3 are enriched in the large intestine^21^. To assess the suitability of our system for studying nutrient uptake, we fed human mini-colons apically with propargyl-choline and then performed a click-reaction to trace its incorporation into the cytoplasm^31^. Notably, we observed a significant uptake and metabolic labeling of phospholipids within the colonic cells of the mini-colons after 12 hours of exposure (**Figure S2A**), suggesting that our system can be leveraged for the study of nutrient and drug uptake, which is cumbersome in organoids.

Consequently, we also demonstrated that our mini-colon system can potentially be used to investigate malfunctions of the epithelial barrier, a common complication in pathological conditions or drug-induced toxicities. While the epithelial barrier is well-maintained under homeostatic conditions (**Figure 1G**), exposure to fungal gliotoxin, which was shown to induce apoptosis in epithelial cells^30^, rapidly compromised barrier integrity by introducing epithelial lesions that allowed the fluorescent probe to leak into the surrounding extracellular matrix (**Figure S2B**). Additionally, human mini-colons offer a valuable platform for simulating intricate microenvironmental interactions that occur *in vivo* but are challenging to replicate using currently available *in vitro* models that lack an extracellular matrix. For establishing these crucial epithelial-stromal interactions, we were able to readily populate the central hydrogel chamber in our hybrid microdevice with intestinal fibroblasts (**Figure S2C**).

We next sought to compare our model to human colon carcinoma Caco-2 cells, widely considered the gold standard for assessing oral drug absorption and ADMETox studies^32^. Even though these cells are of colonic origin and share some features with small intestinal enterocytes, their transformed character and genomic instability raise doubts about their use for basic and translational research. Moreover, Caco-2 cells exhibit aberrant expression of carboxylesterase 1 (CES1) isoenzyme instead of CES2, though levels of CES1 are extremely low in the native human intestine^33^. More importantly, for drug development, cytochrome P450 CYP3A4 is barely expressed in Caco-2 cells, restricting analyses involving CYP3A4-mediated drug metabolism. Genome editing is often used to induce the expression of functional CYP3A4 and CES2 in Caco-2 cells^33–35^. In comparison with this work, qPCR revealed significantly higher CYP3A4 and CES2 expression and the absence of liver-specific CES1 expression in human mini-colons and organoids (**Figure S2D**). Collectively, these early displays of the functionality and significance of human mini-colons set the foundation for future studies on drug permeability and drug-induced gastrointestinal toxicities, for which there is currently no human-relevant *in vitro* system.

### Generating human mini-intestines of ileum origin

We also demonstrate that human ileal stem cells are remarkably conducive to generating perfusable mini-intestines-on-a-chip. Here, we modified the microchannel dimensions to approximate the native ileum crypt dimensions (**Figure S2E**). When seeded into the channel, single intestinal epithelial cells rapidly colonized the hydrogel scaffold and generated an intact, thick epithelium spanning the entire microchannel. Exposing the lumen to a differentiation medium triggered a characteristic epithelial cell turnover, and when coupled with daily perfusion to remove shed cells from the lumen, human mini-guts can be maintained for weeks (**Figures S2E and S2F**). Differentiation conditions also triggered a stereotypical patterning of proliferative (MKI67^+^) and absorptive (ALDOB^+^) cell types in mini-intestines (**Figures S2G and S2H**) to recreate stem/proliferative and differentiated cell zones, which is difficult to achieve with conventional organoids. To investigate cell type composition in human mini-intestines, we compared them to organoids using qPCR, which revealed significantly higher expression of differentiated cell type markers (**Figure S2I**). An immunofluorescence analysis confirmed the presence of secretory cell types with abundant MUC2^+^ Goblet cells and sporadic LYZ^+^ Paneth cells, which are rarely found in classical organoids (**Figures S2J and S2K)**. In stark contrast to human mini-colons, Alcian blue staining of mini-intestines revealed only a thin layer of mucus covering the apical side of the epithelium, consistent with native intestine (**Figure S2L**).

## Discussion

Here we report bioengineered human colon and ileal organoids-on-chip with remarkable cell-type diversity, tissue architecture and function. Our data show that human intestinal stem cells can be coaxed to generate polarized perfusable epithelia with protruding crypt-size invaginations that replicate physiologically relevant architecture of the human gut. Exposing the lumen to a differentiation medium induces a characteristic cell turnover, and repeated perfusion every 24 hours is crucial for eliminating the accumulated shed cells from the lumen. This approach has allowed the reproduction of homeostatic self-renewal and biological longevity and enabled biomimetic patterning of essential intestinal cell types. While recent scRNA-seq analysis have significantly advanced our understanding of human small and large intestine cell type composition^21,23,29^, data on *in vitro*-grown intestinal epithelial tissues are scarce and limited. Efforts have been made to customize culture conditions in order to accommodate enhanced cellular diversity in human small intestinal organoids^3,7^. However, achieving the desired gradients of growth factors and accurately replicating the precise cellular arrangement within these enclosed 3D epithelial cultures has been elusive. Additionally, obtaining simultaneous self-renewal and emergence of fully mature differentiated cell types poses a significant challenge in human colon organoids, due to their strong reliance on Wnt supplementation^4^. Previous bioengineering efforts have generated human colonic epithelia *in vitro*^10–12,14,15^, however the employed differentiation protocols did not yield substantial cellular diversity and maturity or data presented in this regard is notably limited.

In this study we provide a comprehensive overview of the cellular composition and physiological functions of human mini-colons and mini-intestines, which offer a valuable platform for studying patient gut physiology and significantly broaden the real-life applications of intestinal organoids. Drawing on previous findings^4^, we eliminated mitogenic factors on the apical side and provided stem and secretory cell-promoting medium^3^ additionally supplemented with NRG1, which has been shown to increase organoid cellular diversity and maturation ^5,40^, on the basal side. This refined condition drove the concurrent differentiation towards mature colonocyte and secretory cell phenotypes. Indeed, human mini-colons demonstrate the emergence of mature colonocytes, which are spatially localized towards the top of the crypts, while the stem and proliferative cells remain restricted to the bottom. We have also detected the emergence of absorptive BEST4^+^ colonocytes in human mini-colons, which indicates tissue maturation and a close resemblance to the native intestine. This heterogeneity of colonocytes has not been successfully replicated in any *in vitro* system, thus rendering mini-colons a superior model to study nutrient transport and its interconnection to metabolism and onset of different pathologies such as dysbiosis, malabsorption syndromes and inflammatory diseases. Additionally, mini-colons exhibit an exceptional capacity for mucus production by Goblet cells, offering an unprecedented opportunity for investigating the establishment and perturbation of the mucus barrier, which is challenging with currently available systems. The presence of diverse hormone-secreting enteroendocrine cell populations within mini-colons will also facilitate the study of the hormonal signaling system in response to nutrients and pathogens.

Given the luminal accessibility, future studies will focus on establishing bacterial co-cultures in human mini-colons, leveraging the system for drug transport studies, and predicting drug-induced gastrointestinal toxicities. Collectively, we have herein steered the self-organization of human intestinal stem cells into perfusable mini-colons and mini-intestines closely mimicking the native features of the human gut. These models hold significant potential to be leveraged for novel biological and translational applications and will offer valuable insights into longstanding inquiries within a patient-relevant context.

## Limitations of the study

Although human mini-colons capture most of the cellular diversity of the *in vivo* colon, the proportion of Goblet cells appeared significantly lower in mini-colons than *in vivo*, where it represents one of the most abundant cell populations. An immunofluorescence analysis revealed an impressive number of MUC2^+^ Goblet cells, suggesting that this cell type might be particularly challenging to isolate and is more prone to dissociation-induced apoptosis. Furthermore, we did not detect Tuft cells in the mini-colons, suggesting that more refined culture conditions may be required for their enrichment. Also, while we have shown that mini-colons and mini-intestines maintain epithelial self-renewal by observing and removing shed cells on a daily basis, it is necessary to further validate this process using a 5-ethynyl-2’-deoxyuridine (EdU) pulse-chase experiment conducted over several days. This experiment will help confirm whether a complete turnover of the epithelium occurs and provide insights into the turnover rates for both small and large intestinal epithelial cells.

## Supplementary Figures

**Supplementary Figure 1.**
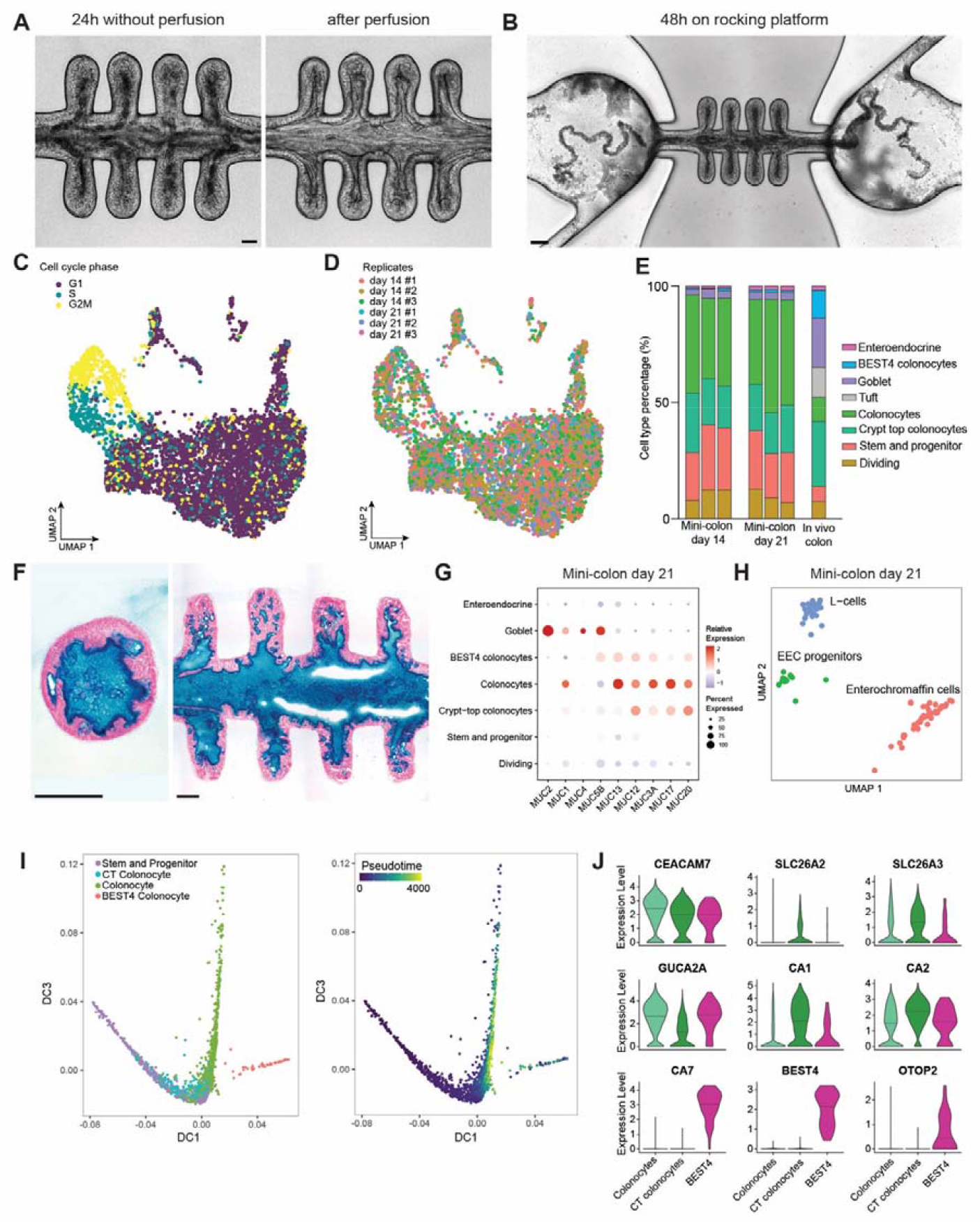
Cell turnover and multilineage differentiation in human mini-colons. **A**, Brightfield image of 15-day-old mini-colon with accumulated dead cells that were shed into the lumen over the course of 24 h without perfusion (left), and following a perfusion pulse (right). **B**, Brightfield image showing a mass of mucus-encapsulated shed dead cell extrusion from human mini-colons cultured on a rocking platform. **C**, Cell cycle states projected onto the UMAP. **D**, UMAP projection of all epithelial cells derived from 3 biological replicates of 14-day-old and 21-day old human mini-colons. **E**, Variability in cell type proportions detected by scRNA-seq among triplicates of 14-day-old and 21-day old human mini-colons compared to the native colon. Average percentages of *in vivo* cell lineages were taken from published dataset on descending colon^23^. **F**, Histological transverse (left) and longitudinal (right) sections stained with Alcian Blue showing lumen of mini-colons filled with mucus in the absence of perfusion. **G**, Dot plot showing the relative expression and the percentage of cells expressing mucin genes across scRNA-seq clusters. **H**, UMAP projection of different Enteroendocrine cell (EEC) subclusters identified in 21-day-old human mini-colons: EEC progenitors expressing NEUROG3, hormone-secreting populations of L-cells (GCG^+^ and PYY^+^) and enterochromaffin cells (TPH1^+^). **I**, Diffusion pseudotime analysis of the colonocyte lineage. Diffusion mapping was performed on mini-colon cells classified as stem and progenitor cells, crypt-top (CT) colonocytes, colonocytes and BEST4^+^ colonocytes. Each dot in the diffusion maps is colored according to its cell type (left) or order in the diffusion pseudotime (right). **J**, Violin plots of the top markers gene expression specific for colonocytes, crypt-top (CT) colonocytes and BEST4^+^ colonocytes. Crossbars indicate median expression. Related to Figure 2.

**Supplementary Figure 2.**
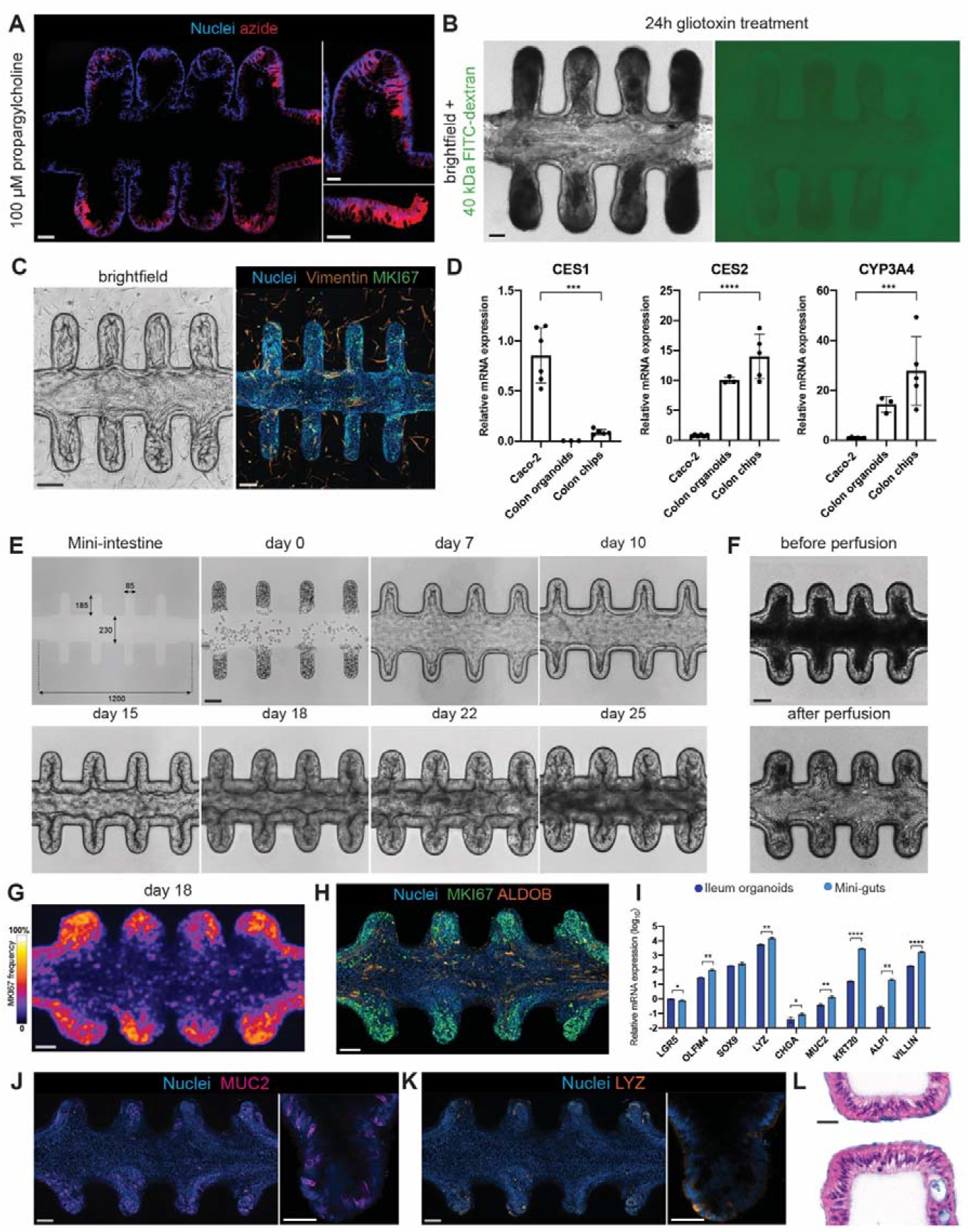
Human mini-colon application potential and mini-intestines derived from ileal stem cells. **A**, Histological cross-section of human mini-colon apically exposed to 100 μM propargylcholine for 24h followed by a click-chemistry reaction to track its absorption by the cells. **B**, Brightfield (left) and fluorescence (right) images of human mini-colons following exposure to gliotoxin show loss of barrier integrity as confirmed by the leakage of 40 kDa FITC-dextran probe into the surrounding ECM. **C**, Potential for modeling epithelial-mesenchymal crosstalk in human mini-colons. Fibroblasts were embedded into the surrounding the tissue ECM to establish co-culture of stromal and epithelial cells. **D**, qPCR gene expression analysis of drug-hydrolyzing enzymes (CES1, CES2) and drug-metabolizing enzyme CYP3A4 in 21-day-old human mini-colons, organoids and 21-day-old Caco-2 cells cultured on Transwell cell culture inserts. Gene expression is normalized to GAPDH and presented as the fold change relative to expression levels in Caco-2 cells.\*\*\*\**P* < 0.0001, \*\*\**P* < 0.001. **E**, Brightfield time-course experiment of epithelium formation in tissue-engineered human mini-intestines established from ileal stem cells. First image depicting dimensions (in μm) of the microchannel approximating *in vivo* ileum crypt architecture. **F**, A 15-day-old mini-intestine with accumulated dead cells that were shed into the lumen over the course of 24 h without perfusion (left), and following a perfusion pulse (right). **G**, Frequency map showing the localization of MKI67^+^ proliferative cells in 18-day-old tissues (right). Average of the maximal intensity projection of a *z*-stack of around 120 μm for *n* = 8 tissues. **H**, Immunofluorescence image of human mini-intestine stained for proliferative cells (MKI67, green) and enterocytes (ALDOB, amber) in 18-day-old tissues (left). **I**, qPCR gene expression analysis of human mini-intestines compared to standard organoids grown in organoid expansion medium (WNRGIFA). Gene expression is normalized to GAPDH and is presented as the fold change relative to LGR5 expression levels in organoids on a log_10_ scale.\*\*\*\**P* < 0.0001, \*\*\**P* < 0.001. **J**, Immunofluorescence image of human mini-intestines stained for Goblet cells (MUC2, magenta). **K**, Immunofluorescence image of human mini-intestines stained for Paneth cells (LYZ, magenta). Scale bars = 100 μm, magnification = 50 μm. **L**, Histological longitudinal cross-sections of 15-day-old mini-intestine tubes stained with Alcian Blue showing thin mucus layer (blue) covering the epithelium and counterstained with Nuclear Fast Red. Scale bar = 25 μm.

**Supplementary Table 1.**
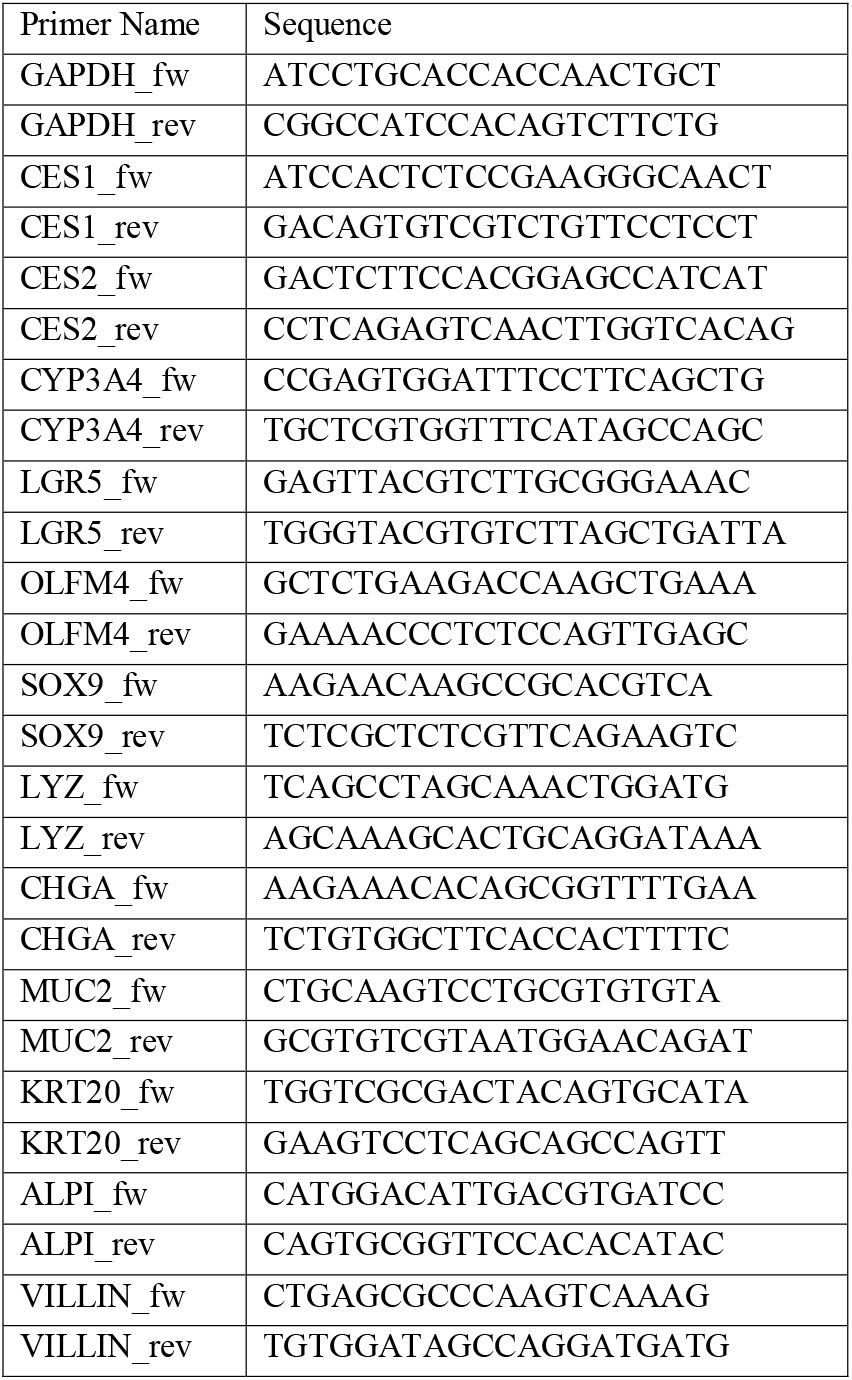
qPCR primer oligos used in study.

## Resource availability

### Lead contact

Further information and requests for resources and reagents should be directed to and will be fulfilled by the lead contact, Matthias Lutolf.

### Materials availability

This study did not generate new unique materials.

## Experimental model and subject details

Healthy human colon and ileum organoids were obtained from the Hubrecht Organoid Technology (HUB) platform (Catalog No. HUB-02-A2-130 and HUB-02-D2-089) and (Catalog No. HUB-14-D2-88 I), respectively. Additional line of human colon organoids used in this study was generously provided by the Schwank lab, University of Zurich, Switzerland.

## Method details

### Human intestinal organoid culture

Human intestinal organoids were cultured as described previously^3^. Organoid expansion medium was composed of Advanced DMEM/F12 with penicillin/streptomycin, 1× Glutamax and 10 mM HEPES, supplemented with 1× B27 supplement (Gibco), 1 μM N-acetylcysteine (Sigma-Aldrich), 0.5 nM Wnt Surrogate-Fc Fusion Protein (U-Protein Express B.V., N001), 100 ng/ml Noggin (EPFL Protein Expression Core Facility), 500 ng/ml R-Spondin 1 (EPFL Protein Expression Core Facility), 100 ng/ml recombinant human IGF-1 (BioLegend), 50 ng/ml recombinant human FGF-2 (Peprotech), 10 nM gastrin, 50 ng/ml recombinant human EGF (R&D, 236-EG) and 500 nM A83-01 (Tocris). Organoids were passaged every 4 days by physical dissociation using fire-polished Pasteur pipettes.

### Caco-2 cell culture

Caco-2 cells were purchased from ATCC (HTB-37) and cultured according to the supplier’s instructions. Briefly, Caco-2 cells were expanded in T75 flasks (Greiner Bio-One, Kremsmunseter, Austria) in complete DMEM medium with penicillin/streptomycin, 1× Glutamax and 10 mM HEPES containing 10% fetal bovine serum (Gibco) at 37°C, 5% CO_2_ and 95% humidity. Cells were passaged at 80% confluence using 0.25% Trypsin EDTA (Fisher). For Transwell culture, Caco-2 cells were seeded at a density of 2.5 × 10^5^ on Transwell cell culture inserts (24-well, Transwell, Costar #3470-Clear, 0.4 μM pore size) and cultured for 21 days in the indicated complete medium. Medium was refreshed every 2–3 days, both 300 μl on apical (insert) and 1000 μl on basal side of the Transwell.

### Culture of human mini-colons

To generate human mini-colon and mini-intestine tubes, organoids were dissociated into single cells in TrypLE express solution (Gibco) containing 2,000 U ml−1 DNaseI (Roche), 1mM N-acetylcysteine (Sigma-Aldrich) and 10 μM Y-27632 (Selleckchem) for 20 min at 37 °C and strained over a 15-μm strainer. Single cell suspension was then introduced through the inlet reservoir of the microfluidic chip and cells were allowed to populate the crypts up to 2/3 of the crypt volume. Non-adherent cells were washed away from the lumen and human mini-guts were subsequently maintained for the first 2 days in full organoid expansion medium with 5 μM Y-27632 but lacking A83-01 and additionally supplemented with 50% fibroblast conditioned medium (1:1). On day 2 fibroblast conditioned medium was reduced to 25% and on day 4 completely removed. Starting from day 7 apical medium was switched to ENR, and basal medium to human organoid expansion medium without EGF and additionally supplemented with 100 ng/ml NRG1 (R&D, 5898-NR) and 500 nM A83-01. Subsequently, medium was changed every other day.

### Immunofluorescence staining

For whole-mount immunostaining human mini-colon and mini-intestine tubes were rinsed with PBS and fixed in 4% paraformaldehyde (PFA) for 30 min at room temperature. After rinsing with PBS, samples were permeabilized with 0.2% Triton X-100 in PBS (2 h, room temperature) and blocked (10% donkey serum in PBS containing 0.01% Triton X-100) overnight at 4 °C. Samples were subsequently incubated overnight at 4 °C with primary antibodies against E-cadherin (1:100, Cell Signalling, 3195T), lysozyme (1:50, Genetex GTX72913), mucin 2 (1:50, Santa Cruz sc-7314), chromogranin-A (1:100, abcam ab15160), Aldolase B (1:100, abcam ab75751), Ki67 (1:50, BD Biosciences, 550609), SLC26A2 (1:100, Proteintech, 27759-1-AP), MUC5B (1:100, Atlas antibodies, HPA008246) diluted in blocking buffer. After washing with PBS for at least 6 h, samples were incubated overnight at 4 °C with secondary antibody Alexa 647 donkey-α-rabbit (1:500 in blocking solution; Invitrogen), Alexa 568 donkey-α-mouse (1:500 in blocking solution; Invitrogen), DAPI (1:5000) and phalloidin-Alexa 488 (1:500; Invitrogen). Following extensive washing for at least 12 hours the samples were imaged on Leica SP8 Inverted microscope using a 25x magnification objective. Bright-field and fluorescent (GFP, FITC-Dextran) images of microfabricated mini-guts and organoids were made on Nikon Eclipse Ti inverted microscope system. Image processing was mainly made in ImageJ (open source). Animated movies were rendered using Premiere Pro (Adobe).

For immunostaining on cryosections a slightly modified protocol was used, where a circle around the tissue was drawn with a hydrophobic SuperHT PAP pen (Biotium) followed by 1 h permeabilization step, 3 h blocking step, overnight incubation with primary antibody, 30 min washing step and 3.5 h incubation with secondary antibody. Washing was performed for 2.5 h in PBS, followed by a mounting step with Fluoromount-G (Thermo Fischer). Glass slides were imaged on Leica SP8 Upright microscope using a 20x magnification objective.

### Microdevice design and fabrication

The microfluidic organ-on-a-chip device was fabricated as previously described^16^. Briefly, the device was composed of three main compartments: a hydrogel compartment for organoid culture in the center, two open media reservoirs flanking the hydrogel compartment, and inlet/outlet open media reservoirs for perfusion of the microchannel. Matrix chamber was connected to a pair of inlets for media perfusion and an extra inlet through which the matrix was loaded. Inlet and outlet reservoirs were designed to function as air-bubble traps and allow for the connection of tubings and facilitate injection of medium in small quantities (functional tests, bacterial co-culture). The platform for microfluidic chips was fabricated using conventional soft-lithography methods and poly(dimethylsiloxane) (PDMS) molding at the Centre for Micronanotechnology, EPFL. The resulting PDMS chips were briefly exposed to oxygen plasma and irreversibly bonded onto a glass substrate. Chips were sterilised with UV and kept sterile until further use.

### Hydrogel loading and microchannel fabrication

An ECM solution containing 75% (v/v) neutralized 6 mg/ml bovine TeloCol®-6 Type I Collagen Solution (Advanced Biomatrix) and 25% (v/v) Matrigel was injected into the matrix compartment of the microdevice through the matrix loading inlet and incubated at 37°C for 3 min, after which the cell loading inlets and media reservoirs were filled with human BMGF media. Generation of microtracks in hydrogel was performed using a nanosecond laser system (1 ns pulses, 100 Hz frequency, 355 nm; PALM MicroBeam laser microdissection system (Zeiss)) equipped with a 10X/0.25NA objective, at a constant stage speed and a laser power. A pattern of consecutive parallel lines was created in Wolfram Mathematica and then imported into the PALM MicroBeam system’s interface. Pattern was positioned along the microdevice matrix compartment, covering its entire length, at Z=220 μm from the glass surface. Laser power and etching speed were adjusted to achieve 110-120 μm high microtrack in hydrogel. Following microtrack generation, microchannels were perfused with base medium and microdevices were maintained at 37°C in 5% CO_2_ humidified air.

### Histology

The samples were fixed in 4% PFA for 40 min at room temperature and washed 3 times in 1× PBS. To extract a block of hydrogel containing mini-gut tubes from the microfluidic chips hydrogel compartment was cut around its perimeter using a razor blade, which was subsequently transferred to an Eppendorf tube and subjected to consecutive overnight incubations in 1) 30% sucrose, 2) 30% sucrose with Cryomatrix (Epredia) mix (1:1) and 3) 100% Cryomatrix. Finally, specimens were placed in a histology mold filled with Cryomatrix and frozen on dry ice. Sectioning was done using a Leica cryostat CM3050S at −20 °C. Section thickness was set at 16 μm. For staining, sections were hydrated in distilled water and immersed in Alcian Blue pH 2.5 solution for 25 min, counterstained with Nuclear Fast Red, dehydrated and mounted with a xylene-based glue. Hematoxylin and eosin staining was performed at the EPFL Histology Core Facility using a Ventana Discovery Ultra automated slide preparation system (Roche). Sections were imaged on a LEICA DM 5500 microscope, DMC 2900 colour camera. Image processing was performed using ImageJ with standard contrast and intensity level adjustments.

### Epithelial barrier integrity experiments

Medium in the apical perfusion channel was replaced by medium containing 1 mg/ml FITC-dextran (40 kDa, Sigma, FD40). To evaluate barrier integrity imaging was performed at different time-points: 0, 2, 12, 24 h. To compromise barrier integrity in human mini-colons, fungal toxin gliotoxin from Gliocladium fimbriatum (Sigma, G9893) was used at a final concentration 13 μg/ml (1:1000) in apical medium.

### Propargyl-choline nutrient uptake experiments

Human mini-colons were apically exposed to propargylcholine (Aobious, AOBT7378) at a final concentration 100 μM in apical medium for 12 h. Subsequently, samples were fixed in PFA and cryosections were prepared as described above. Glass slides with cryosections were then stained via a click-chemistry reaction as previously described^41^: by incubating for 10–30 min with 100 mM Tris (pH 8.5), 100 μM CuSO4, 100 μM fluorescent azide Alexa Fluor 546 (from 10 mM stock in DMSO), and 100 mM ascorbic acid (added last to the mix from a 0.5 M stock in water). The staining mix was prepared fresh each time and was used for staining tissue cryosections immediately after addition of ascorbate. After the staining, the slides were washed several times with 0.2% Triton X-100 in PBS, mounted with Fluoromount-G (Thermo Fischer) and imaged as described above.

### Fibroblast conditioned medium preparation

Human Intestinal Myofibroblasts were purchased from Lonza (CC-2902) and cultured in SmGMTM-2 Smooth Muscle Cell Growth Medium-2 (CC-3182) according to manufacturer’s instructions. For conditioned medium preparation, myofibroblasts were used between passage 2-4 and expanded in T150 flasks until 100% confluency. Flasks were then washed once with PBS and incubated 3 times in DMEM medium with 100 μg ml−1 penicillin– streptomycin, 1× Glutamax and 10 mM HEPES (Gibco) exchanging the medium every 30 min to eliminate any traces of serum. Medium was then changed for the 4^th^ time and cultures were maintained for 48 h until collection. Conditioned medium was then centrifuged at 1500 rpm for 5 min and concentrated 30x times in Amicon® Ultra-4 Centrifugal Filter Unit with 30 kDa MWCO cut-off (Merck Millipore). Concentrated conditioned medium was then filtered using 0.2 μm filters and frozen until use.

### Real-time qPCR

21-day-old cultures of human mini-colons or mini-intestines and Caco-2 cells maintained on Transwell cell culture inserts were briefly washed with PBS and lysed with RLT buffer (500 μl) from the RNeasy Micro Kit (Qiagen) supplemented with dithiothreitol (40 mM, Sigma). The RNA was then extracted using RNeasy Micro Kit (Qiagen). The complementary DNA was synthesized by using iScript cDNA synthesis kit (Bio-Rad) at 0.5 μg (20 μl) per sample on a UnoCycler (WVR). The qPCR was performed with Power SYBR Green PCR Maxter Mix (Applied Biosystems) on a QuantStudio 6 (Applied Biosystems, QuantStudio Real-Time PCR software v.1.3). The primer sequences used can be found in the Supplementary Table 1. The relative gene expression levels were normalized to GAPDH and further standardized as indicated in the figure legends.

## Quantification and statistical analysis

### Epithelium coverage quantification

To quantify the epithelium formation efficiency bright-field images of human mini-colons were taken every day with the same acquisition parameteres. Machine learning software ilastik was used to classify pixels into three classes (background, epithelium-covered area, epithelium-non-covered area) and to generate probability maps. Probability maps were then imported into Fiji and the epithelium area was quantified using custom-made macro. For batch processing trained ilastik project file was generated and a pixel classification prediction was performed using ilastik plugin in Fiji.

### Single-cell RNA-sequencing data processing

14-day-old and 21-day-old human mini-colons in biological triplicates were extracted from the chips and dissociated to single cells using 4 mg/mL Protease VIII (Sigma) in dPBS containing 10 μM Y-27632 on ice for approximately 45 min with trituration using a P1000 micropipette every 10 minutes. Following one washing step in PBS+2% BSA (Gibco), cell suspensions were incubated for 20 min on ice with TotalSeq™-C anti-human Hashtag oligos (HTOs) #1-6 in PBS+2% BSA (1:500, Biolegend, product codes: 394661, 394663, 394665, 394667, 394669, 394671 were used). Cells from each condition and replicate were washed three times with PBS+2% BSA, pooled together and filtered through low-volume 10 μm cell strainers (PluriSelect). All cell suspensions were recounted to achieve a uniform concentration of 2000 cells per microliter before pooling for 10× capture. The cell hashing and cDNA libraries were constructed using 10x Genomics Chromium Next GEM Single Cell 5’ Reagent Kits v2 (Dual Index) reagents and sequenced using Illumina protocol using NovaSeq 6000 reagents, with around 50,000 reads per cell. The reads were aligned to GRCh38 with Cell Ranger v.6.1.2. Raw count matrices were imported in Seurat 4.3.0 using RStudio 2022.12.0+353 and single live cells were selected on the basis of the number of detected genes (approximately 2,500–5,000) and fraction of mitochondrial genes (around 0.05–0.15). Demultiplexing of cells to their original sample-of-origin and identification of cross-sample doublets was performed using *HTODemux* function. Suspected doublets, debris and contaminants were excluded. The number of cells after filtering was 2050 (day 14 mini-colons) and 5115 (day 21 mini-colons). The data were normalized to 10,000 counts per cell, and log1p-transformed using the natural logarithm (referred to as log(expression)). The datasets were aligned using *harmony* package before dimensionality reduction with UMAP. Graph-based Louvain clustering (resolution of 1) using the *FindClusters* function of Seurat yielded 7 clusters that were assigned to known cell types by manual examination of cluster markers and expression of canonical cell type markers. The cell cycle phase of each cell was inferred using the *CellCycleScoring* function in Seurat. Visual representations of the data were generated using Seurat internal functions or ggplot2 and cosmetic adjustments were made in Adobe Illustrator.

### Diffusion Pseudotime Analysis

For diffusion pseudotime analysis, we used scRNA-seq data of Stem and progenitor, crypt-top (CT) colonocytes and late colonocytes in 21-day-old mini-colons. Diffusion mapping analysis was performed using the *DiffusionMap* function in the R bioconductor package *destiny* (version 3.14.0)^42^ with the following parameters: sigma = ‘local’, k = 500. Diffusion pseudotime from the tip of the stem cell branch was obtained using the *DPT* function in the *destiny* package^42^.

## Data and code availability

The single-cell RNA-sequencing data have been deposited in the Gene Expression Omnibus (GEO), accession number GSE235072. The code required to reproduce the single-cell RNA-sequencing analysis detailed above will be available in the github repository upon request or manuscript acceptance. Any additional information required to reanalyze the data reported in this paper is available from the lead contact upon request.

## Supplemental information

**Supplementary Video 1 – Establishment of epithelial barrier and cell turnover in human mini-colons**.

**Supplementary Video 2 – Mucus secretion in human mini-colons**.

## Author contributions

O.M. and M.P.L. conceived the study and wrote the manuscript. N.B. analyzed raw scRNA-seq data. O.M. designed experiments and analyzed data. M.N. designed and microfabricated the microdevice.

## Acknowledgments

We thank J. Langer for generating laser microtrack patterns, A.Chrisnandy for the help with propargylcholine click-chemistry reaction, L. Francisco Lorenzo and N. Gjorevski for helpful discussions. We acknowledge valuable support from the following EPFL core facilities: CMi, Histology, Gene Expression. This work was funded by support from the Swiss National Science Foundation (SNSF) research grant 310030_179447, the EU Horizon 2020 Project INTENS (#668294-2) and Ecole Polytechnique Fédérale de Lausanne (EPFL).

